# Swift Formation of Optimal Single Spheroids towards *In-Vitro* 3-Dimensional Tumour Models

**DOI:** 10.1101/2021.12.16.472941

**Authors:** Kinana Habra, Joshua R.D. Pearson, Stéphanie E. B. McArdle

**Affiliations:** Chemistry department, School of Science and Technology, Nottingham Trent University, Nottingham NG11 8NS, UK; John van Geest Cancer Research Centre, School of Science and Technology, Nottingham Trent University, Nottingham NG11 8NS, UK; Centre for Health, Ageing and Understanding Disease, School of Science and Technology, Nottingham Trent University, Nottingham, UK

**Keywords:** carnosine, glioblastoma, prostate cancer, 3-D model, single spheroid, high throughput, U87 MG, SEBTA-027, SF188, DU145, TRAMP-C1, IncuCyte

## Abstract

Monolayer cell culture, while useful for basic *in vitro* studies, are not physiologically relevant. Spheroids, on the other hand provide a more complex 3-dimensional (3D) structure which more resemble the *in vivo* tumour growth thereby allowing results obtained with those on proliferation, cell death, differentiation, metabolism, and various anti-tumour therapies to be more predictive of *in vivo* outcomes. However, the cost associated with their generation often involve expensive, plate, media, and growth supplements, which have limited their use for high throughput experiments. The protocol herein presents a novel and rapid generation for single spheroids of various cancer cell lines, U87 MG; SEBTA-027; SF188, brain cancer cells, DU-145, TRAMP-C1, prostate cancer cells, in 96-round bottom well plates. Cells are washed with anti-adherent solution, and the homogeneous compact spheroid morphology was evidenced as early as 24 hours after 10 minutes centrifugation for the seeded cells. By using confocal microscopy, the proliferating cells were traced in the rim and the dead cells were found inside the core region of the spheroid. The H&E stain of spheroid slices and the western blotting were utilised to investigate the tightness of the cell packaging by adhesion proteins. Carnosine was used as an example of treatment for U87 single spheroids. The protocol allows the rapid generation of spheroids, which will help towards reducing the number of tests performed on animals.

## Introduction

Glioblastoma multiforme (GBM) is the most malignant primary form of glioma with prognosis of around a year after diagnosis (Jiang and Uhrbom, 2012; Ostrom et al., 2018). The complete removal of GBM tumour is impossible to be achieved by surgery alone due to the complexity of the tumour cells’ structure and their migration away from the bulk of the tumour to the neighbouring brain parenchyma (Lara-Velazquez et al., 2017). Moreover, the tumours are frequently resistant to radiation and chemotherapy (Bao et al., 2006; Messaoudi et al., 2015). Despite the availability of various anti-cancer treatments, the treatment of solid tumours remains challenging, this is due to the tumour tissues developing three-dimensional (3D) solid morphology with cell-cell and cell-matrix contacts. Due to the multicellular resistance, actual tumours are more resistant to therapy than the monolayers of cell cultures (Trédan et al., 2007). Single spheroids of tumour cells mimic the biological properties of micro-metastases and vessel distal regions of tumours by retaining the architectural, morphological, and physiological characteristics of the tumour. Thus, there is a serious need to look towards new effective therapeutic strategies to cure this terminal disease (Cuzzubbo et al., 2021).

*In vitro* spheroid models bridge the complexity gap between monolayer and *in vivo* tumour growth because spheroids exhibit many of the tumour microenvironment features by the avascular region of tumour tissues which resist drug diffusion (Ivascu and Kubbies, 2006). The limitation in the delivery of anticancer treatments to the far situated cancer cells from the tumour blood vessels occurs due to the major characteristics of solid tumours which include cell to cell adhesion, extracellular matrix, large distances between blood vessels, and high interstitial fluid pressure. Thus, modelling the multicellular resistance of tumour cells may contribute to developing *in vitro* findings into effective clinical therapies (Phung et al., 2011). Whenever animal studies are unfeasible, 3D spheroids are able, to some extent, mimic the clinical environment. Therefore, *in vitro* spheroid models are the most commonly used tool to assess drug penetration (Hjelstuen et al., 2009).

We have previously demonstrated that carnosine can be safely used as a potential therapy for GBM, however, the model of sustained release of the active substrate requires further investigation (Habra et al., 2021). The *in vitro* applications for spheroids are numerous for cancer cell lines. However, forming compact round spheroids in 3D was an obstacle (Phung et al., 2011). To overcome this problem, generating single spheroids has been developed using various culture techniques (Mueller-Klieser, 2000). However, many limitations remain regarding the standardisation of the spheroids’ high throughput production, homogeneity, and cost-effectiveness of these 3D models (Kelm et al., 2003; Kunz-Schughart et al., 2016; Santini and Rainaldi, 1999). Here, we describe a detailed protocol to establish an *in vitro* 3D single spheroid model which can be utilised to identify potential new therapeutic approaches. This cost-effective protocol is reliable and be applied to investigate *in vitro* studies of various tumour targeting drugs, antibodies and immunoconjugates (Winter et al., 2003). Here we describe the time of cell attachment when the cells stop shrinking and start growing as a spheroid by monitoring the diameter or bright field area over time, the presence of cell death, using stains that enable the detection of the dead/ live cells, and we visualise the inner and outer area of the spheroids to examine the invasion. Eventually, the H & E histology test and the protein expression for the tightness of the generated spheroids using the proposed protocol.

## Materials and methods

### Cell Culture

The human glioblastoma U87 MG-Red-FLuc (Bioware Brite, PerkinElmer, Waltham, Massachusetts, USA) was the authorized cell line in all experiments. These cells were incubated in Opti-MEM Reduced Serum Medium (Gibco™, Thermo Fisher Scientific, Waltham, Massachusetts, USA) culture medium, supplemented with fatal bovine serum up to 10%. The antibiotic puromycin (Gibco®, Thermo Fisher Scientific Waltham, Massachusetts, USA) was added after the initial thaw at 2 µg/mL. The incubation was at 37°C in a humidified atmosphere containing 5% CO_2_. Other cell lines were cultured in their specific media by following the same protocol outlines. The human Glioblastoma SEBTA-027 (Recurrent GBM cell line derived from the right parieto-occipital region of a 59-year-old female) and SF188 (GBM cell line derived from and 8-year-old male) were cultured in Gibco™ DMEM, high glucose, GlutaMAX™ Supplement, and 10% fetal calf serum (Gibco™, Thermo Fisher Scientific, Waltham, Massachusetts, USA). Both cell lines were a generous gift from the University of Portsmouth, neuro-oncology group. The human Prostate cancer DU145, HTB-81™ (American Type Culture Collection ATCC, Virginia, USA) were cultured in Eagle’s minimum essential medium modified to contain Earle’s Balanced Salt Solution, non-essential amino acids, 2 mM L-glutamine, 1 mM sodium pyruvate (BioWhittaker® Medium EMEM Cell Culture Media, Lonza, Maryland, USA), and 10% fetal calf serum. The murine prostate cancer TRAMP-C1 (C57Bl/6 mice cells which are derived from prostate adenocarcinoma cells from TRAMP mice) were cultured in Dulbecco’s Modified Eagle Medium 4.5 g/L glucose w/L-Gln w/ sodium pyruvate (DMEM, Lonza, Maryland, USA), and 10% fetal calf serum. This cell line was provided by Matteo Bellone (University of Milan, Milan, Italy). The human breast cancer BT-549 are ductal carcinoma (American Type Culture Collection ATCC, Virginia, USA) were cultured in Corning RPMI 1640 Medium with L-Glutamine (Corning™ RPMI 1640 Medium, New York, USA), 10% fetal calf serum and 0.023U/mL insulin. The murine breast adenocarcinoma Py230 (American Type Culture Collection ATCC, Virginia, USA) was cultured in Corning Medium F-12K with L-glutamine (BioWhittaker® Medium F12K Medium, Lonza, Maryland, USA), and 10% fetal calf serum, 0.1% MITO+ serum extender (Corning®, New York, USA).

### Generation of spheroids

A volume of 50 µl of Anti-Adherence Rinsing Solution (STEMCELL Technologies, Cambridge, UK) was added to each well of a clear green coded 96-well round base plate for suspension (Sarstedt, Nümbrecht, Germany). After 15 minutes all wells were washed with 50 µl of serum free media. U87 MG cells were grown as a monolayer then detached with Trypsin (Sigma-Aldrich, St. Louis, Missouri, USA) to generate a single-cell suspension. The cells were seeded (400 cells/well) in 100 µl full media. Directly, the spheroid formation was initiated by centrifuging the plates at 3700 RPM for 10 min. The plates were incubated under standard cell culture conditions at 37° C, 5% CO_2_ in humidified incubators. The full media had been replaced each other day.

### Localization of cell death and proliferation within spheroids

U87 MG spheroids were generated from seeding 400 cells per well in 100 µl of media into a 96-well, round-bottom plate as explained previously. The spheroids were stained by IncuCyte^®^ Cytotox red for counting dead cells (250 nM, Essen Bioscience). The spheroids were transferred to the IncuCyte S3 Live Cell Analysis System (Essen Bioscience Inc., Ann Arbor, Michigan, USA), images were snapped with 4× objective lenses in each well every hour inside an incubator over 10 days. The localisation of the red dead cells within the spheroids was assessed by phase-contrast images using the red channel to evaluate the real-time cell membrane integrity and cell death. The total phase and the red fluorescent areas with mask were quantified for different days using IncuCyte^®^ spheroid analysis software (Version. 2020B, Essen Bioscience Inc., Ann Arbor, Michigan, USA).

### Imaging and microscopy studies

The single spheroids were stained following the instructions of each kit. The dead cells were stained by IncuCyte^®^ Cytotox red for counting dead cells (250 nM, Essen Bioscience) during the generation process. The cyanine nucleic acid dye permeated cells with compromised cell membranes. The green dye CFSE Cell Division Tracker Kit (BioLegend, San Diego, California, USA) and DAPI (Sigma-Aldrich, St. Louis, Missouri, USA) were used to stain the live cells and the nucleus within before taking pictures. The morphology of the spheroids was assessed and recorded using the IncuCyte S3 Live Cell Analysis System and confocal laser scanning microscope (Leica, Wetzlar, Germany) by a 5× objective using the following settings: sequential scanning, ex/em: Mitotracker: 543/599 nm, Hoechst: 405/461 nm. The size of the spheroids was analysed using IncuCyte® spheroid analysis software (Version. 2020B, Essen Bioscience Inc., Ann Arbor, Michigan, USA), and ImageJ software (version. 1.44, National Institute of Mental Health, Bethesda, Maryland, USA).

### Hematoxylin and eosin H & E stain of spheroid slices

U87 MG spheroids were generated from 400 cells. Prior staining was cultivated for 1, 3, 5, 7, 10 days, washed once in PBS, fixed with 10% formalin (Merck, Darmstadt, Germany). The sparoids were washed in PBS and transferred to disposable biopsy Molds to encapsulate them with Epredia™ HistoGel™ Specimen Processing Gel (Fisher-Scientific, UK). Each gel capsule was moved to a cassette and loaded inside a tissue processor (Excelsior AS, Thermo Scientific, Germany). The program was set to start with 6 times of alcohol, then 3 times of xylene, 3 times of wax. Each cycle time was 10 minutes. Then the capsules were embedded in paraffin using (Histostar, Thermo Scientific, Germany) with a temperature range of -3 °C to -12 °C. Microtone (Leica, RM2235) sections of 5 μm were placed on Super Frost glass slides (Menzel-Glaser, Thermo science, Germany) and allowed to dry for 2 hours at 37° C. The sections were deparaffinized by 2 changes of xylene for 5 min, rehydrated by 2 changes of 100% ethanol, followed by washing in 70% ethanol for 1 min. After a short single rinse in distilled water, the sections were stained for 20 min in Mayer’s hematoxylin (Merck) and placed for 20 min under running tap water, then dipped in 1% Scott’s Tap. The slides were observed under the microscope. If the nuclei were too blue, a quick dip in Acid alcohol followed by a tap water rinse. If it was not blue enough the staining was repeated from hematoxylin step. Sections were counterstained with eosin (Merck) for 2 min, rinse quickly in tap water, dehydrated by a dip in 70% ethanol, followed by 2 changes in 100% ethanol for 2 min each, and 2 changes of xylene for 2min. The slides were mounted in DPX (Sigma-Aldrich, Germany), and left to air dry for 2 hours. spheroid sections were assessed by bright field microscopy.

### Immunoblotting of cell membrane proteins

A confluent between 70% to 80% U87 MG monolayer culture served as positive control. After 24 hours and 10 days, the spheroids were pooled from one 96-well round bottom plate. All spheroids were washed twice with ice-cold PBS, then lysed in 500 µl RIPA lysis buffer (50 mM Tris-HCl pH 8, 150 nM sodium chloride, 0.1% sodium dodecyl sulfate (SDS), 0.5% sodium deoxycholate, 1% Triton x100, 1mM EDTA) containing protease inhibitor cocktail (Sigma). The lysates were vigorously vortexed and placed on ice every 10 minutes for a 30-minute period. After vortexing samples were centrifuged for 15 min at 15,000 rpm at a temperature of 4°C. The protein concentration was determined by a BCA protein assay (Sigma). A total of 50 µg protein extract was mixed with 5× Laemmli loading buffer (50% glycerol, 10% SDS, 0.25% bromophenol blue, 250 mM Tris-HCl pH 6.8, 5% β-mercaptoethanol) resolved on a 10% SDS-PAGE gel and a wet transfer was performed with 25 mM Tris, 192 mM glycine and 20% methanol for 90 minutes at 100 V onto a polyvinylidene fluoride (PVDF) membrane. Membranes were then blocked with a 5% milk in TBST solution and then probed with primary antibodies directed against β-actin (Sigma), vinculin (Abcam), vimentin and E cadherin (Cell Signaling Technologies). After incubation with primary antibodies, HRP-linked secondary antibodies (Cell Signaling Technologies) were used to detect bound primary antibodies in combination with Clarity Western ECL substrate (BioRad Laboratories).

## Results and discussion

The IncuCyte results in Figure 1 show the ability of the U87 MG cells to form a single spheroid after around 24 h. Utilising carnosine in a sustained release designed experiment we demonstrated the shrinking effect on the single spheroids and compared it to the untreated spheroid control. The change in morphology of the single spheroids was monitored using the IncuCyte spheroids software for a period of 7 days to create (Video 1) which proved the penetration of carnosine to the live cells in the single spheroids. Carnosine was added every 48 hours over the 7 days to mimic the sustained release therapy, the series of carnosine concentrations reflected the critical amount to hinder the viability of the spheroid which was above 100 mM.

**Fig 1.**
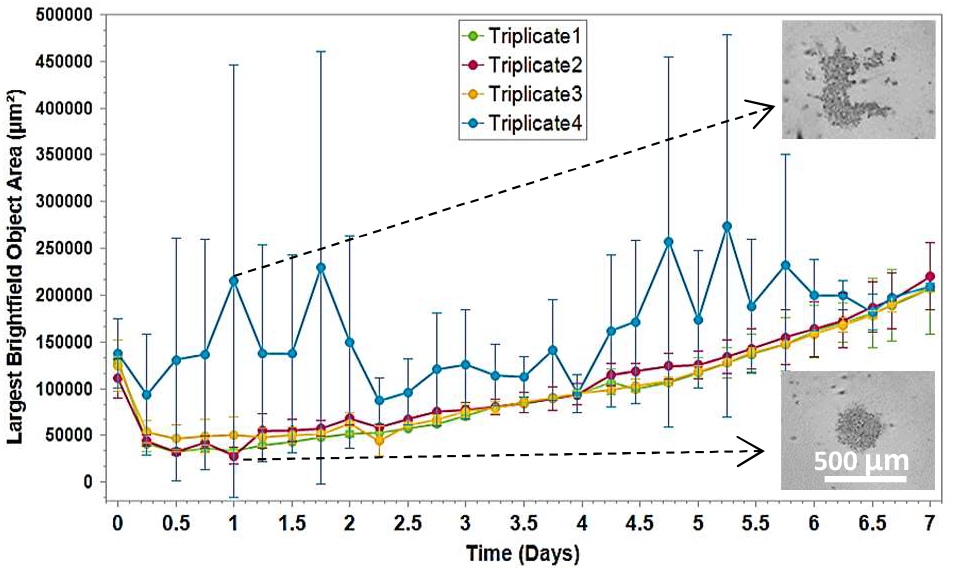
IncuCyte S3 live cell system (4×) Live spheroids images analysis shows the proliferation curves of the confluence ratio of U87 single spheroid upon using the protocol steps. The first triplicate failed to convert the aggregates to spheroids without using the washing solution. However, following the protocol showed consistent reproducibility for the single spheroids on three triplicates 2, 3, and 4. The images show the difference between the shape of the aggregates and the successful shape of the spheroid.

The critical step in the protocol is the use of the anti-adherence washing solution which consists of an amphipathic component to prevent cell adhesion (Schlenoff, 2014). We found that using 96 well round bottom plates promoted the formation of homogenous spheroids instead of aggregates (Fig 1). The method was applied horizontally and vertically in the plate to monitor the consistency and reproducibility in all locations. The IncuCyte live cells system showed the 3D structure of the spheroids which was tightened after the same period in all seeded wells. All spheroids were grown consistently over a period of 12 days (Fig 2).

**Fig 2.**
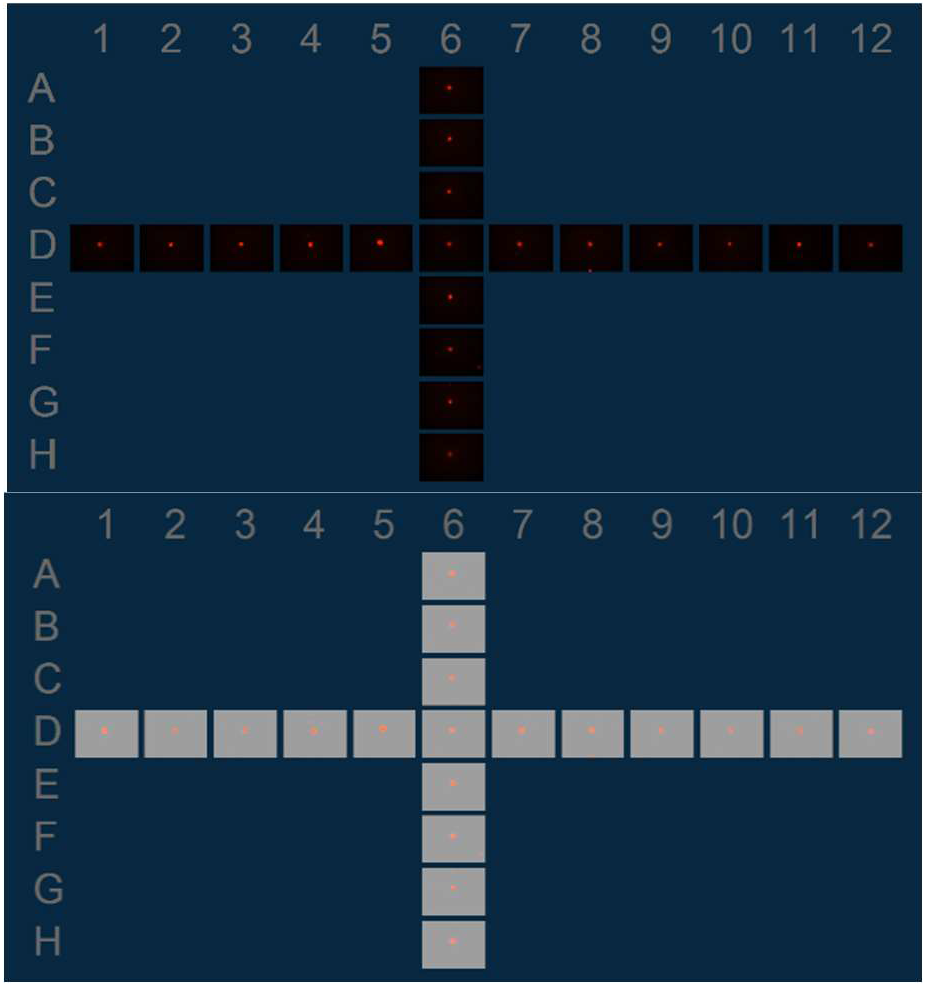
IncuCyte S3 live cell system (4×) Live spheroids images plate images show the homogeneous single spheroids in all wells locations (day 3). The Cytotox red stain was used to label the dead cells in the centre of each spheroid. The images show the similarity in all spheroids when detecting the red channel with and without the phase.

Spheroid morphology with cells in suspension well plates versus other types of round bottom 96 well plates revealed the overall transformation from forming aggregates to generating tight morphology of single spheroids where the dead cells (in red) were localised in the centre. The confocal microscopic images showed the live cells (green with blue nucleus) proliferated outwards, the volume of the single spheroid was significantly increased after 5 and 10 days (Fig 3).

**Fig 3.**
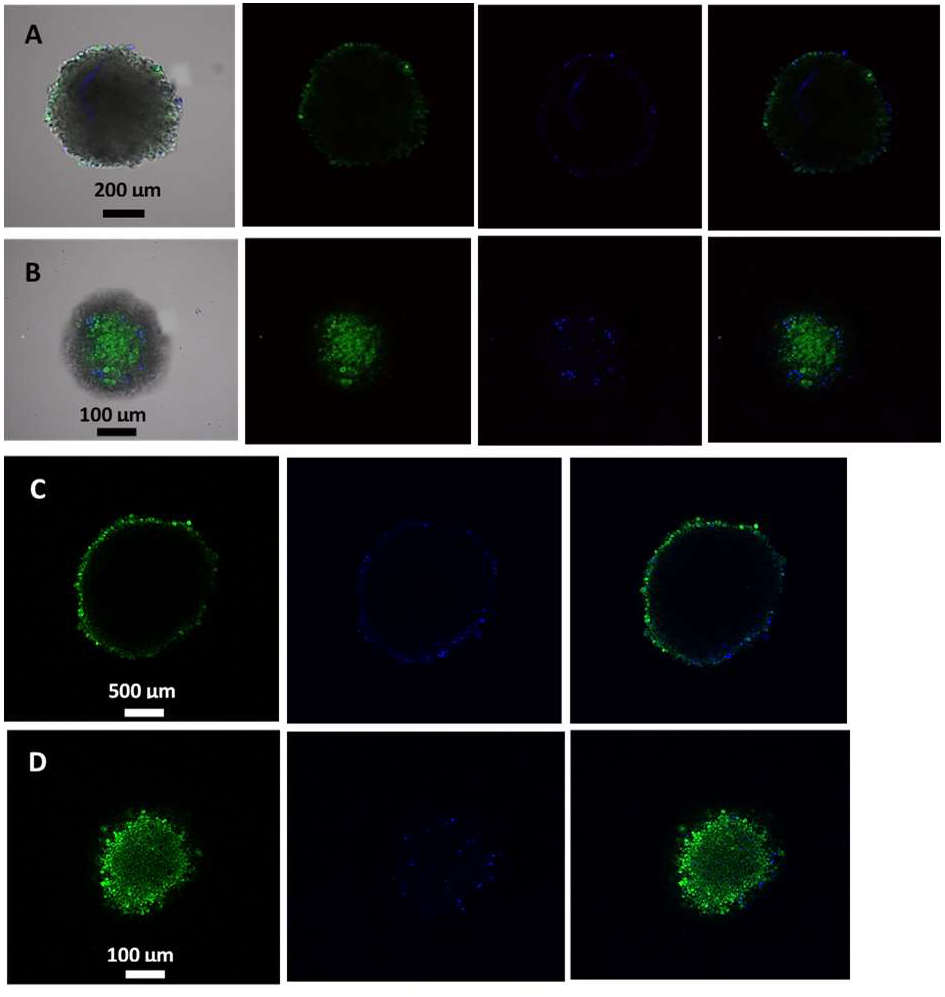
Confocal images show the distribution of the green live cells with the blue nucleus which are located close to the rim of the spheroid in day 5 and 10. **(A, C)** The 5- and 10-days spheroid images consequently across the centre show the dark shade were the dead cells are existed. **(B, D)** The 5- and 10-days spheroid images consequently across the rim show the tightness of the cells surrounding the spheroid from the edges.

To trace the localisation of the dead and live cells, an IncuCyte image was taken using the green channel filter to show the live CFSE stained cells surrounding the Cytotox red colour taken up by dead cells at the core of the spheroid (Fig 4, A-D). This method has the advantage of assessing the mobility of the tumour cells by the invasion study of the U87 MG cells from the surface of the spheroid. The invasion mask can be applied, and the area of the dead cells can be subtracted from the whole spheroid (Fig 4, E).

**Fig 4.**
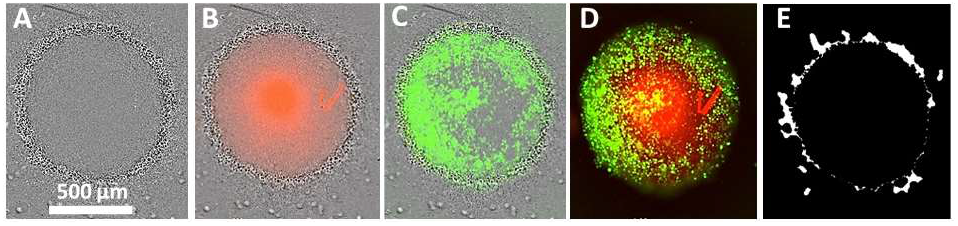
IncuCyte S3 live cell system (4×) Live spheroids images after 10 days. **(A)** The phase image shows the confluence of the live cells around the spheroid distributed uniformly. **(B)** The phase with the red channel filter image shows the localisation of the dead cells in the centre of the spheroid. **(C)** The phase with the green channel filter image shows the 3D localization of the live cells around the spheroid. **(D)** The overlaps of the red and green filter show the 3D shape of the cells dead/ live in the same spheroid. **(E)** The masks of the whole and invasion of the cells which could be used to monitor the effect of any applied treatment over time and convert it to a curve using the IncuCyte spheroid invasion software.

To evaluate the homogeneity and the dynamic of the growing single-well spheroids, the increase in sizes was quantified by the IncuCyte spheroids software. Unlike aggregates, the spheroids were regular in shape, displayed a uniform round geometry, and exhibited a narrow size variation. The spheroids size was estimated using the mean confluence of the phase area for 4 spheroids. The cells were gathered over the first 24 hours which was displayed as a shrinkage in the detected area. However. After a day and 4 hours all wells showed homogeneous spherical single spheroid from U87 MG cell line. The diameter of the U87 MG spheroids formed from 400 cells per well ranged from 216 ± 9 µm after 24 hours to 475 ± 8 µm after 5 days, then 847 ± 11 µm after 10 days. The homogeneity of the spheroids was reflected by the small standard deviation ranging of the mean (Fig 5, A). Using a different type of plate resulted in a complete failure in the production of single spheroid with or without washing the wells with the anti-adherence solution. Without using the washing solution, the cells started to adhere at the bottom of the plate. Even with washing the wells, the images showed an incomplete circle. Thus, the cells were not able to gather properly during the centrifugation (Fig 5, B & C).

**Fig 5.**
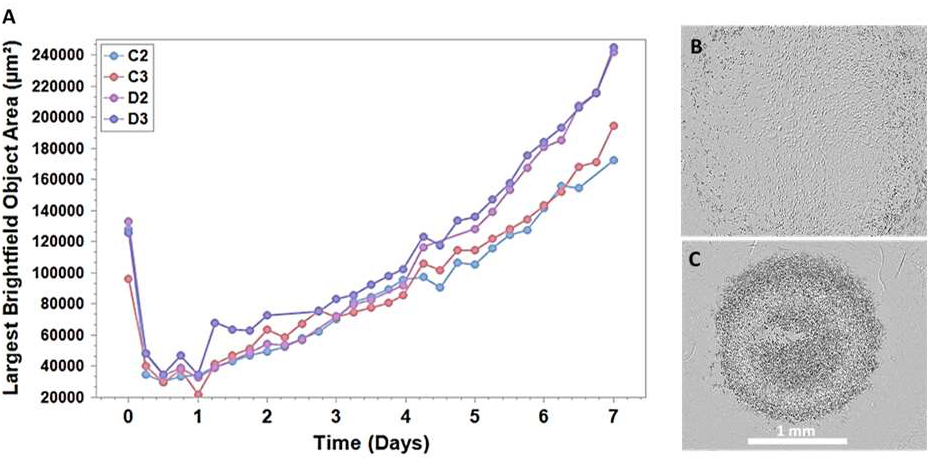
IncuCyte S3 live cell system (4×) Live spheroids images analysis shows **(A)** the proliferation curves of the confluence ratio of U87 single spheroid for four spheroids shrinked to the half size after 24 hours then grew over time. The images show the negative effect of using different type of plate. **(B)** the image shows the instant adherence of the cells when using different plate without washing. **(C)** The image shows the cells gathered when the anti-adherence solution. However, the plate material was not suitable to allow the tightness of the cells. A crack in the middle appeared after the centrifugation directly.

To investigate the tightness of the cell packaging in the single spheroid, histological sections of spheroids grown for 1, 3, 5, 7, 10 days in culture were examined. H & E is the combination of hematoxylin which stains cell nuclei with a purplish-blue colour, while eosin stains the extracellular matrix and cytoplasm pink. Figure 6 shows an example of H & E stain from the centre and the rim section of a spheroid at day 7. This evidenced that the density of U87 MG cells was high in the core region, whereas the daughter cells around the rim gathered to tighten and increase the spheroid size. The figure displayed an H & E stain of the middle section of the spheroid which encompassed the complete tight structure of the spheroid from core to rim. The rim of the spheroid consisted of evenly layers of packed cells toward the centre despite the death of the cells.

**Fig 6.**
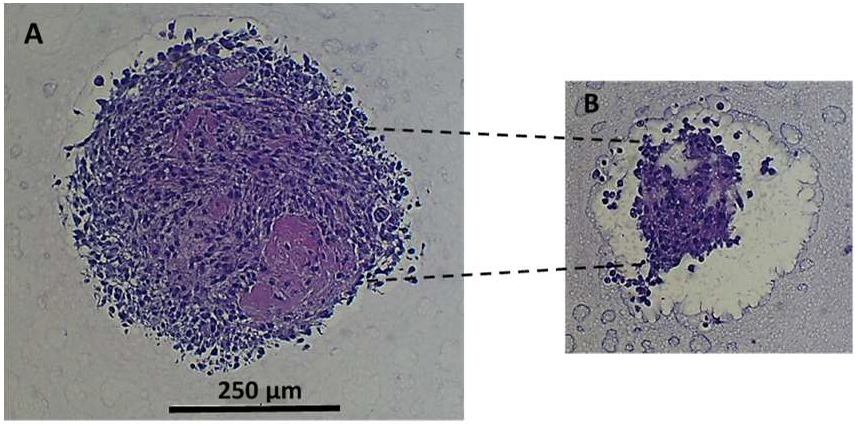
Cell tightness and interaction analysis of U87 MG spheroids. H & E stain of spheroid slices from **(A)** the core area, and **(B)** top rim area of 7 days spheroid generated from 400 cells.

In this project, we have generated single-well spheroids from 5 different tumour cell lines starting from same cell density per well (Fig 7). To optimize the mimicking of different stages of avascular tumour regions, the size of the spheroids can be adjusted by seeding different cell numbers prior to centrifugation. The applications which target primarily viable cells at the rim and core region of the spheroids should be small, whereas large spheroids will harbour a larger centre with a necrotic area (Carlsson et al., 1983). The resulted single spheroids could be transferred by ease of mechanical access to any plate or cell culture vessel for further investigations or analysis. Cell viability in the spheroid might be significantly affected differently by the 3D microenvironment of multicellular contacts such as the expression of tight junction molecules, which establish a delay the shrinking in size. On average the spheroids reach a stable symmetrical size with smooth surfaces after 24 to 48 hours before growth started. However, some cell lines will require a longer time to adjust before starting to grow such as DU145 which needed around 5 days to produce firm spheroid (Fig 7).

**Fig 7.**
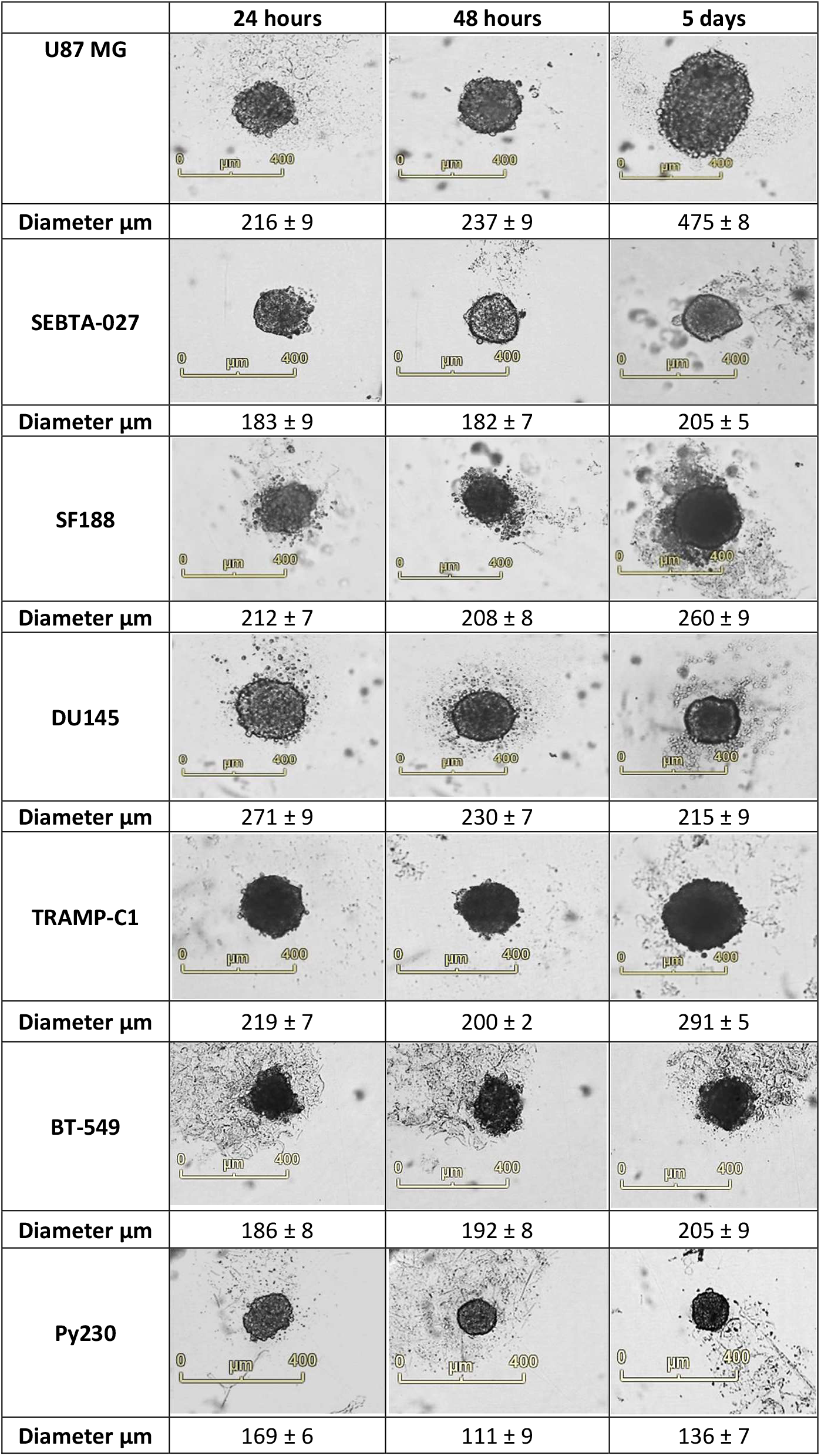
IncuCyte S3 live cell system (4×) Live spheroids images show the potential of using the protocol to obtain single spheroids from various cell lines. The comparison shows consistency in the mean size and similar standard deviation for 6 individual samples from each cell line (U87 MG, SF188, SEBTA-027 human brain tumours, DU145 prostate human tumour, and TRAMP-C1 prostate murine tumour). The scale bar is 400 µm.

During spheroid formation, a small proportion of cells did not integrate inside the sphere and lost cell-cell adhesion properties. The reason for this separation is the gravity-sedimentation (Stadler et al., 2018). However, adhesion and tight junction proteins are the principal factors involved in turning cell aggregates into spheroids. Western blotting revealed a down regulation of E-cadherin and vimentin in spheroid cultures compared to cells grown as a monolayer (Fig 8). These findings imply that these spheroids adopt a highly invasive mesenchymal phenotype as opposed to an epithelial phenotype (Iwadate, 2016). This epithelial to mesenchymal transition (EMT) has previously been identified in spheroids formed using the CAL33 head and neck squamous cell carcinoma (Essid et al., 2018). These EMT changes are indicative of a cancer stem cell phenotype; a population known to be highly treatment resistant and responsible for tumour recurrence (Lathia et al., 2015). As a result of these findings spheroids represent an excellent *in vitro* model for the testing potential therapies and provide a platform for *in vivo* tumour formation.

**Fig 8.**
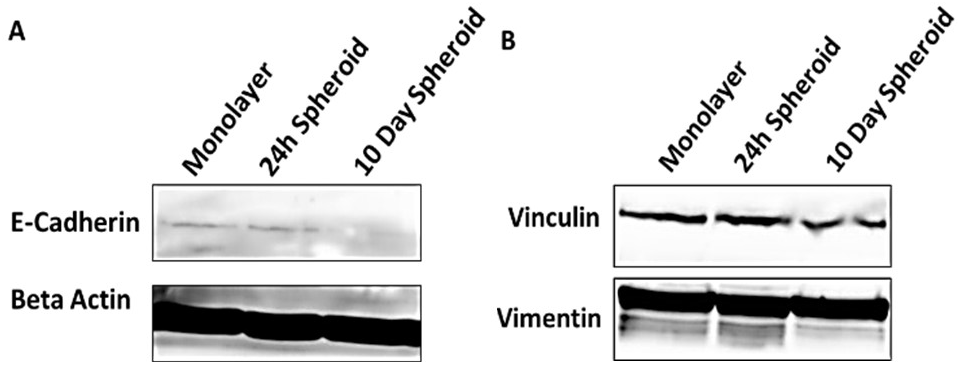
Expression of **(A)** E-cadherin and Beta Actin. **(B)** Vinculin and vimentin in U87MG cells grown either in a monolayer, or as spheroids derived from 400 cells for 24 hours and 10 days.

## Conclusion

Monolayer cultured tumour cells exhibit less resistance to therapeutic interventions than *in vivo* cells. Developing a 3D model that represents the tumour microenvironment is important, especially when trying to bridge the gap between *in vitro* and *in vivo* tumour models. This protocol produced uniform single spheroids without any additives. Consistently, this method demonstrated reproducible high throughput U87 MG spheroids formation within a reasonable timeframe and cost. The spheroids exhibited the ability of being solid and allowed the penetration of the carnosine treatment, simultaneously. Examination of the U87-MG-Red-Fluc spheroids also revealed that these spheroids mimic tumour morphology, with a necrotic core of dead cells being visible much like the necrotic core often observed in GBM tumour histology. Optimising different cell lines will provide the research workers with an easily accessible and realistic model for investigating the effectiveness of various cancer therapies *in vitro*.

## Author Contributions

K.H. performed the experiments and wrote the first draft. J.R.D.P. performed the Western blot experiment and contributed to the final manuscript. S.E.B.M. contributed to the final manuscript. All authors have read and agreed to the published version of the manuscript.

## Data Availability Statement

The data presented in this study are available on request from the corresponding author.

## Acknowledgments

The authors wish to acknowledge Graham Hickman for his technical support with the electron microscopy studies. The authors appreciate the efforts of Gareth Williams in the histology work. K.H. would like to thank the NTU & CARA fellowship program for the generous award and support.

---

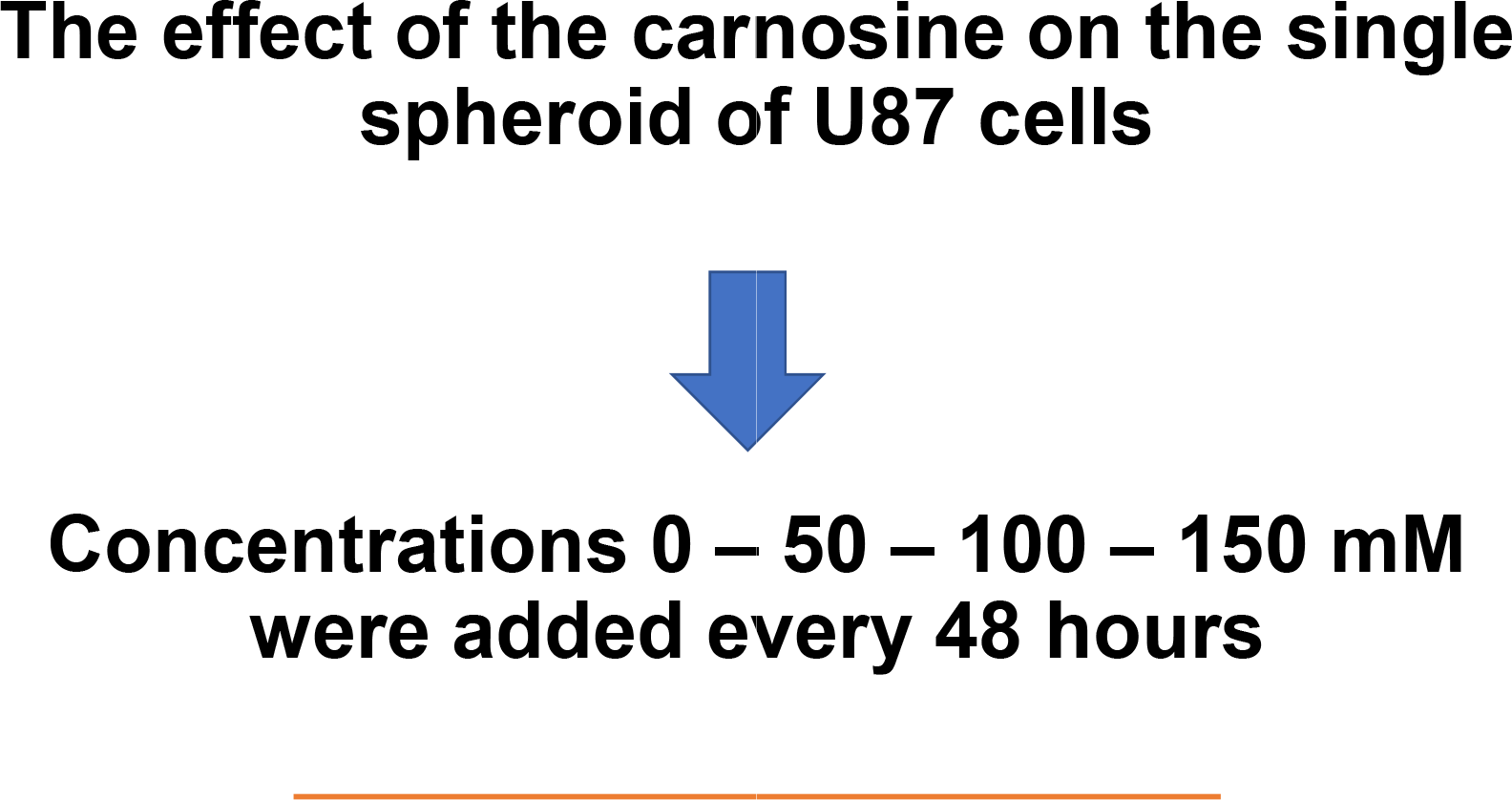

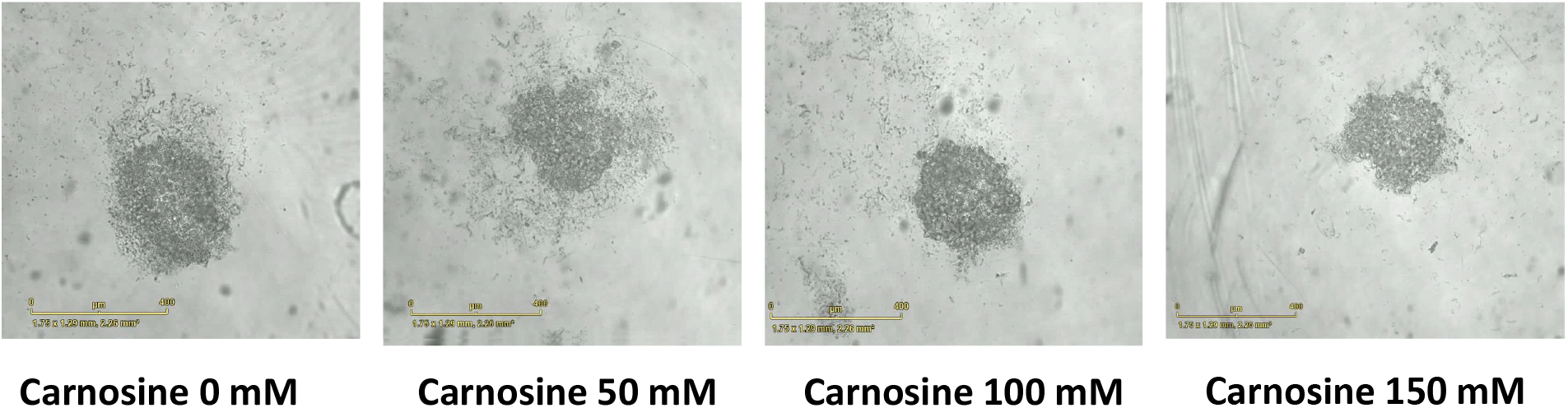

